# Dug: A Semantic Search Engine Leveraging Peer-Reviewed Knowledge to Span Biomedical Data Repositories

**DOI:** 10.1101/2021.07.07.451461

**Authors:** Alexander M. Waldrop, John B. Cheadle, Kira Bradford, Alexander Preiss, Robert Chew, Jonathan R. Holt, Nathan Braswell, Matt Watson, Andrew Crerar, Chris M. Ball, Yaphet Kebede, Carl Schreep, PJ Linebaugh, Hannah Hiles, Rebecca Boyles, Chris Bizon, Ashok Krishnamurthy, Steve Cox

## Abstract

**Motivation:** As the number of public data resources continues to proliferate, identifying relevant datasets across heterogenous repositories is becoming critical to answering scientific questions. To help researchers navigate this data landscape, we developed Dug: a semantic search tool for biomedical datasets utilizing evidence-based relationships from curated knowledge graphs to find relevant datasets and explain *why* those results are returned.

**Results:** Developed through the National Heart, Lung, and Blood Institute’s (NHLBI) BioData Catalyst ecosystem, Dug has indexed more than 15,911 study variables from public datasets. On a manually curated search dataset, Dug’s total recall (total relevant results/total results) of 0.79 outperformed default Elasticsearch’s total recall of 0.76. When using synonyms or related concepts as search queries, Dug (0.36) far outperformed Elasticsearch (0.14) in terms of total recall with no significant loss in the precision of its top results.

**Availability and Implementation:** Dug is freely available at https://github.com/helxplatform/dug. An example Dug deployment is also available for use at https://search.biodatacatalyst.renci.org/.

## Introduction

The ability to interrogate large-scale data resources is becoming a central focus of many research efforts. The U.S. National Institutes of Health (NIH) and other public funding agencies have supported data generation at unprecedented scales through projects such as Trans-Omics for Precision Medicine (TOPMed) (University of Washington Department of Biostatistics, 2020), All of Us (The “All of Us” Research Program, 2019), and Helping to End Addiction Long-Term (HEAL) (Collins *et al.*, 2018). From these efforts, the ability to integrate data within and across disjoint and complex public data repositories is quickly replacing data scarcity as a primary bottleneck to research progress. While successful data integration efforts have resulted in novel diagnostics, therapies, and prevention strategies, researchers often lack necessary tools for navigating this complex data landscape (Powell, 2021).

There is a growing need for comprehensive search tools that identify datasets relevant to a researcher’s particular scientific question. Despite recent NIH emphasis on making research data more Findable, Accessible, Interoperable, and Re-Usable (“FAIR” data principles) (Wilkinson *et al.*, 2016), the diversity of public data repositories has proven to be a formidable barrier to developing intelligent search strategies. To illustrate, consider that the NIH alone currently refers data submission to more than 95 domain-specific repositories (NIH Data Sharing Resources, 2020). Often, more established repositories like the NIH database of Genotypes and Phenotypes (dbGaP; https://www.ncbi.nlm.nih.gov/gap/) require studies to submit only free-text descriptions of experimental variables. The complexity of “heart attack” vs. “myocardial infarction” exemplifies the challenges of identifying relevant datasets among the growing corpus of nonstandardized biomedical datasets.

Emerging techniques for Natural Language Processing (NLP) are enabling semantic search over biomedical datasets. We define semantic search as search that considers the intent and context of the query as opposed to a purely lexical approach (Tran *et al.*, 2007). Many existing search and annotation tools successfully employ methods for named entity recognition and disambiguation to annotate free text with synonyms or similar ontology terms (Bell *et al.*, 2019; Chen *et al.*, 2018; Canakoglu *et al.*, 2019; Huang *et al.*, 2016; Laulederkind *et al.*, 2012; Pang *et al.*, 2015; Soto *et al.*, 2019; Chapman *et al.*, 2020).

Similar NLP techniques have been successfully employed by semantic-aware dataset search engines like Google Dataset Search (Brickley *et al.*, 2019) and Datamed (Bell *et al.*, 2019) to make data more discoverable. Broadly, both tools index study-level metadata describing high-level features of each dataset (e.g., study description, abstract) to increase findability. For Datamed, study metadata are augmented with semantic concepts identified by NLP annotation tools to increase findability. Google Datasets extends this concept by mapping study metadata elements to Google’s internal knowledge graph, used to augment metadata with additional relevant search terms. For a more complete review of dataset search tools, see Chapman et al. (2020).

Despite the utility of these tools, there remains a need for a biomedical dataset search engine that can empower true data re-use. A critical shortcoming of the search engines described above is a focus on *study-level* metadata to the exclusion of *variable-level* metadata (i.e., the features measured as part of this study); two or more studies can be combined and integrated with only a single conceptually overlapping variable, enabling a broader set of possible data integrations.

There also remains a need for a biomedical dataset search engine that combines context-awareness with domain specificity. The ability to show second and third order connections can be helpful for data discovery but requires a knowledge graph describing how biological entities and phenomena are related. Despite the practical utility of Google’s proprietary knowledge graph for general search, the provenance, depth, and quality of its biomedically relevant connections are not easily verifiable. There remains a need for a search tool capable of leveraging evidencebased biological connections to show researchers datasets useful for hypothesis generation or scientific support.

Therefore, we present Dug (https://github.com/helxplatform/dug): a semantic search tool for biomedical datasets that leverages NLP technologies and ontological knowledge graphs curated by biomedical subject matter experts to intelligently identify datasets relevant to a user query. Dug is designed specifically for data integration and re-use by allowing users to search over variable-level metadata and high-level study metadata. Dug is also the first biomedical search engine that can explain *why* it returns what it returns (Fig. 1). For example, if a search for “cancer” were to return a dataset that measured “asbestos exposure,” Dug can show the user—often with links to supporting literature—that this variable was returned due to the causal linkage between asbestos exposure and certain types of cancer. Here, we discuss Dug’s motivations, architecture, functionality, and evaluation, and demonstrate its successful deployment in the NHLBI’s BioData Catalyst Ecosystem (National Heart Lung and Blood Institute *et al.*, 2020).

**Fig 1:**
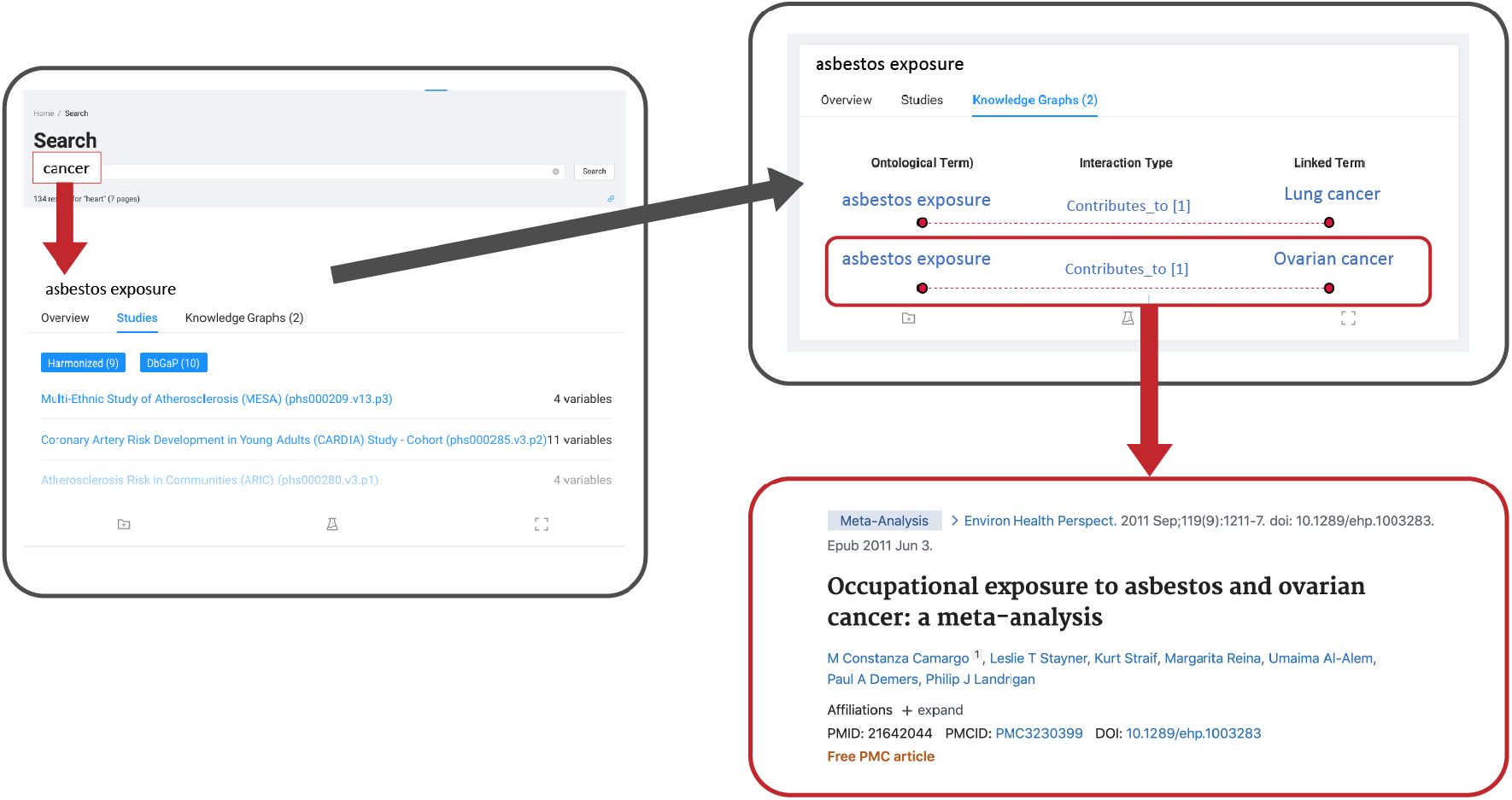
The Dug web portal leverages knowledge graph connections with supporting links to PubMed literature to explain why certain results are relevant to a user’s query.

## Implementation/Methods

Dug is designed to find relevant datasets for a query like “lung cancer,” and to allow users to discover datasets they would not have found using lexical or even synonym-based search engines. Here, we discuss the computational architecture underpinning this functionality.

Dug consists of two components: A **Dug API service** that orchestrates metadata ingestion, indexing, and search, and the **Dug search web portal** that displays results to end users (Fig. 2). The ingestion/indexing pipeline is designed to:

1. Parse heterogenous study metadata formats into a common Dug metadata format.
2. Automatically annotate free-text descriptions of study variables with a set of ontological identifiers.
3. Expand annotations with relevant terms returned from knowledge-graph queries.
4. Index each study variable, associated ontological concepts, and set of knowledge graph answers to an Elasticsearch (Kuć and Rogozinski, 2016) endpoint.

**Fig. 2:**
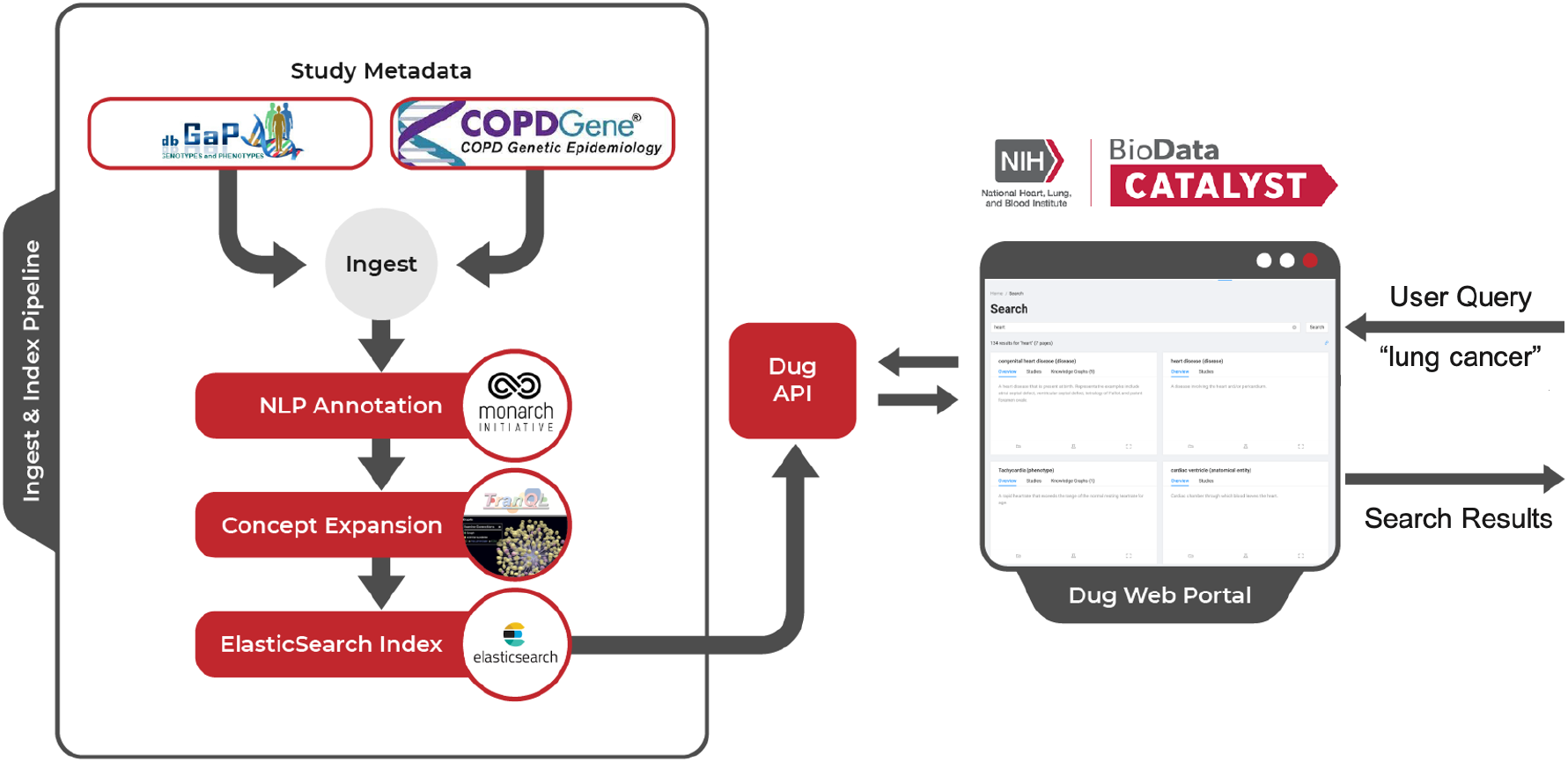
High-Level Dug Architecture. Dug makes study metadata searchable by parsing heterogenous metadata formats into a common format (ingest), annotating metadata using NLP tools to extract ontology identifiers from prose text (NLP annotation), searching for relevant connections in federated knowledge graphs using Translator Query Language (TranQL) (concept expansion), and indexing this information into an Elasticsearch index. Dug’s web portal utilizes a flexible API to query and display search results back to end users.

Finally, the Dug search portal is a web-based application that leverages the Dug API to display search results and render explanatory knowledge graphs. Below, we discuss each component in detail.

### Ingestion and Indexing Pipeline

#### Data ingestion

Dug ingests and indexes study *metadata* (e.g., text descriptions of study variables) as opposed to actual study data. To accommodate the diversity of metadata formats available across public data repositories, our ingestion pipeline abstracts out retrieval modes (e.g., local file, network file) and data parsing formats (e.g., XML, JSON). Similar to the Data Tags Suite (DATS) metadata schema (Sansone *et al.*, 2017), Dug parses diverse metadata formats into a common *DugElement* metadata model, which defines a standard set of metadata required for indexing. Dug can be flexibly adapted to ingest nearly any metadata format by extending its plug-in interface to parse input data into *DugElement* objects. Supplemental materials include a detailed discussion of the *DugElement* metadata model.

#### Data Annotation

Dug’s data annotation module extracts a set of biomedical ontology identifiers from ingested metadata elements using tools for named entity recognition (Fig. 3). Typically, we use the `/nlp/annotate` endpoint exposed by the Monarch Initiative’s (Mungall *et al.*, 2016) Biolink API. The underlying Biolink model provides a high-level data model for representing biomedical knowledge (Reese *et al.*, 2021) and can be used to integrate across domain-specific ontologies. Monarch’s API service accepts prose text as input and returns a set of ontological identifiers with additional information in JSON.

**Fig. 3:**
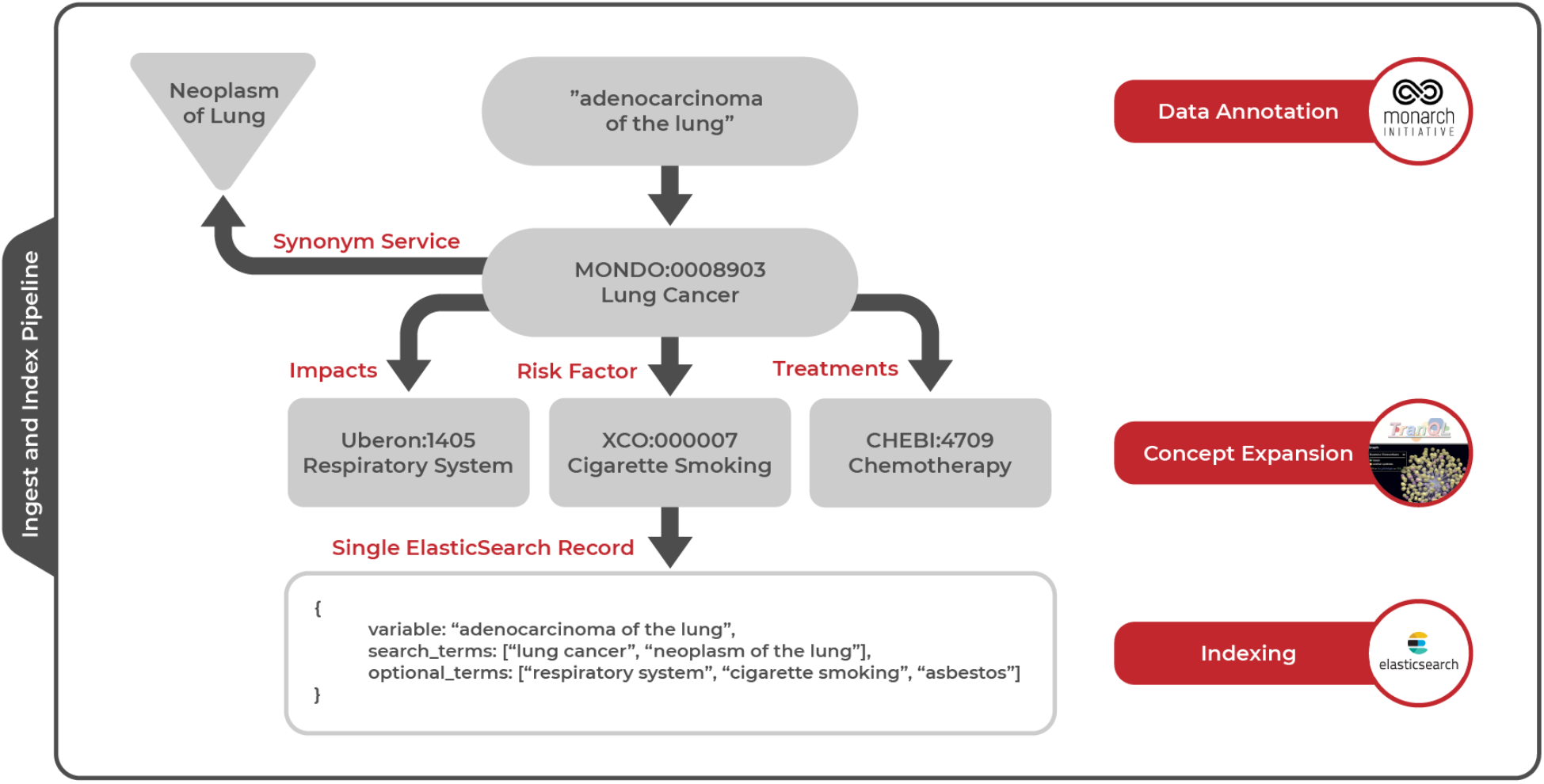
Detailed example of ingest and index pipeline. After ingesting a variable called “adenocarcinoma of the lung” from study metadata, Dug uses NLP methods to annotate the variable with an ontology identifier for “Lung Cancer” from the Mondo disease ontology. The resulting identifier is used to gather synonyms for lung cancer such as “Neoplasm of lung” from an external API service. During concept expansion, Dug leverages TranQL to query knowledge graphs for other ontological concepts related to lung cancer through certain predicates; in the figure, we are looking for risk factors, treatments, and anatomical entities impacted by lung cancer. During indexing, all terms discovered through annotation and concept expansion are combined with the original metadata into a single Elasticsearch record so that queries against any of these terms will yield the initial variable measuring “adenocarcinoma of the lung.”

To accommodate NLP services and tools for biomedical named entity recognition, Dug also abstracts out the Annotation module. To extend Dug’s annotation interface, developers can create a child class specifying the new API endpoint and define a function converting successful API responses into an internal data structure called a *DugIdentifier*, which defines a minimal set of ontological information needed for downstream processing (e.g., id, name, Biolink type).

By converting free text to standardized ontology identifiers, we can leverage semantic web services supporting this nomenclature to gather additional information about each identifier. Dug utilizes a normalization service (necessary to interoperate with Biolink knowledge graphs, discussed below) to transform identifiers to the preferred equivalents (https://github.com/TranslatorSRI/NodeNormalization). Additionally, Dug uses an ontology metadata service for fetching identifier names, descriptions, synonyms, and Biolink types (https://onto.renci.org/apidocs/).

#### Concept Expansion

Dug’s ability to retrieve contextualized search results and explain these connections to end users is undergirded by a process we call **concept expansion**, which further annotates ontological identifiers by identifying relevant connections within ontological knowledge graphs (Fig. 3).

The data structure for concept expansion is a knowledge graph, with nodes representing entity types (e.g., disease, gene, chemical exposure) and edges providing predicates that describe the relationship between entities; Biolink predicates include “causes,” “is associated with,” and “is expressed in.” Biolink provides an “upper ontology” that defines connections across domain specific ontologies (e.g., Mondo, ChEBI, HP). For example, the ChEBI “asbestos” identifier might be linked to the Mondo “lung cancer” identifier via the Biolink “risk_factor” predicate. Inclusion/exclusion of specific ontologies, and creation of links across ontologies, is curated by subject matter experts as part of the Biolink initiative.

To contextualize metadata within a knowledge graph, we leverage data integration approaches developed through the NCATS Biomedical Data Translator (Biomedical Data Translator Consortium, 2019). Chief among these are ROBOKOP (Bizon *et al.*, 2019) and TranQL (Translator Query Language; https://tranql.renci.org) (Cox *et al.*, 2020).

Dug leverages TranQL to gather an expanded set of ontological concepts related to those extracted via NLP annotation. Dug allows platform administrators to define the set of TranQL query templates used to retrieve related ontology identifiers. Below is an example of a query template used to retrieve chemical risk factors for a disease:

~~~
FIND Chemical_Entity -> Risk_Factors -> Disease WHERE
Disease == {Query Ontology ID}
~~~

During concept expansion, Dug uses templates to substitute actual ontological identifiers from the previous NLP annotation step to retrieve a set of relevant terms for a specific variable. In Fig. 4, Dug returns a list of chemical risk factors for lung cancer by substituting an ontology identifier for lung cancer (MONDO:0008903) into the query template. The actual query becomes:

~~~
FIND Chemical_Entity -> Risk_Factors -> Disease WHERE
Disease == MONDO:0008903
~~~

**Figure 4:**
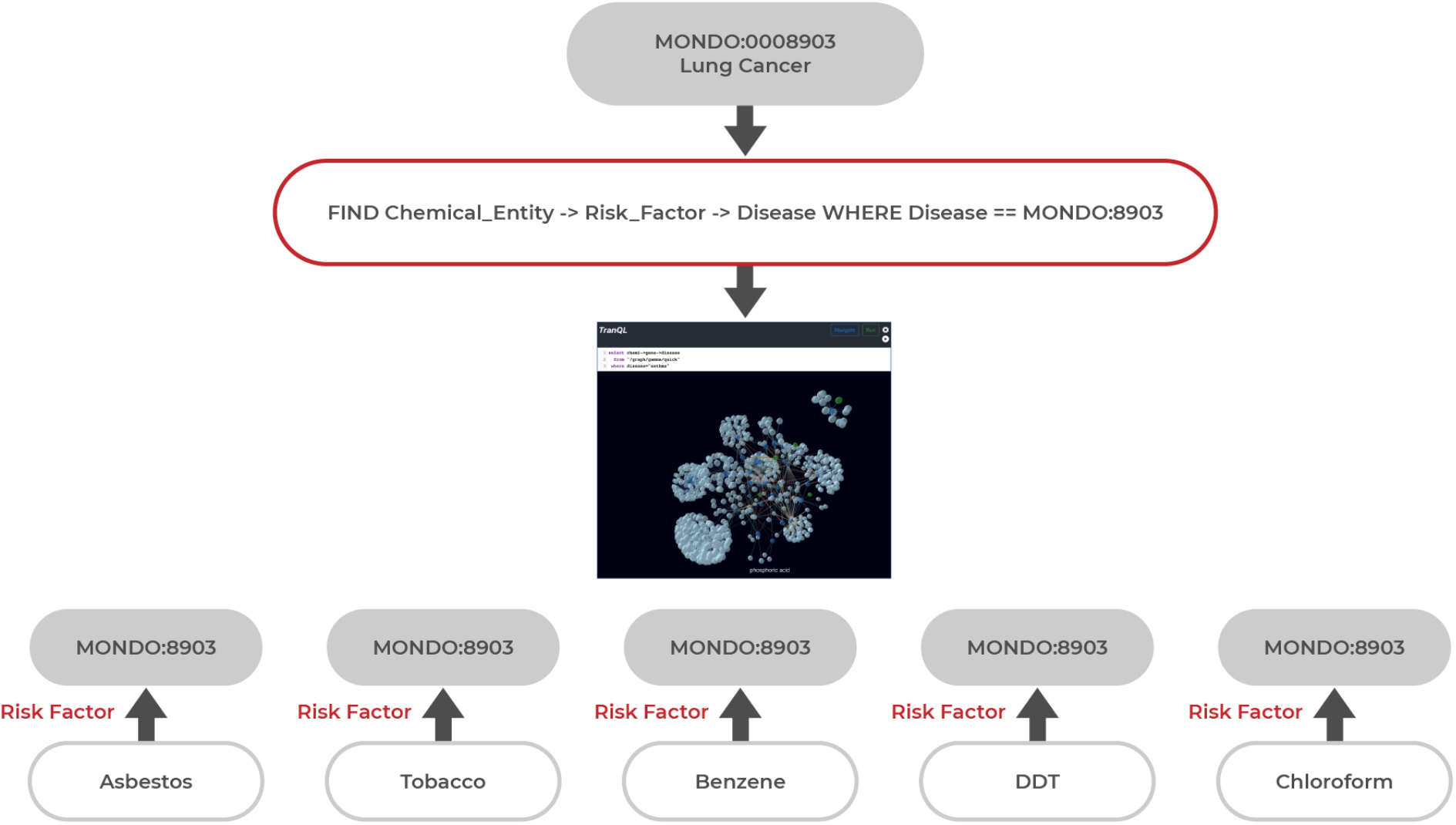
An example TranQL query for chemical risk factors of lung cancer. Each TranQL answer returned is a knowledge subgraph linking one chemical element node back to the query node.

The “answers” returned by TranQL queries are used to increase the search relevance of related concepts and to provide a basis for including explanations for links that led to the result (e.g., peer reviewed literature supporting knowledge graph links). TranQL “answers” are also knowledge graphs: a set of nodes (ontological identifiers), edges (predicates), and metadata including names, descriptions, synonyms of ontological identifiers, and PubMed literature links supporting each edge (Fig. 4).

#### Data Indexing

After annotation and concept expansion, the resulting data structure must be indexed. To maximize speed and flexibility of Dug’s search functionality, Dug’s back-end search architecture is implemented as a set of linked Elasticsearch indices. Elasticsearch is a stack of flexible components that allow users to quickly index and search over large volumes of structured text data. Elasticsearch provides the basic functionality to “save” structured bits of text called “documents” into databases called “indices” that can be queried by downstream users. By default, Elasticsearch uses a “fuzzy” term frequency-inverse document frequency (TF-IDF) scoring algorithm for text retrieval, a purely lexical search that allows some character mismatches and ranks importance of results by the inverse frequency of each token in the complete set of documents.

Dug’s semantic capabilities result from combining ingested study and variable metadata with terms harvested through annotation and concept expansion into a single Elasticsearch record (see Fig. 3). For each metadata variable, Dug’s indexer adds a “search_terms” field to the original record containing the names and synonyms of each identifier added via data annotation. The indexer adds an “optional_terms” field by traversing the names of knowledge graph nodes added during concept expansion. Dug’s results *always* include text matches from the original metadata, since Dug’s Elasticsearch query searches over the original metadata fields parsed during ingestion *and* the expanded fields added during annotation and concept expansion.

Dug quickly organizes search results by higher-level concepts through partitioning 1) ingested metadata records, 2) core ontological concepts, and 3) expanded knowledge graph answers respectively into three separate Elasticsearch indices. Any ontological identifier extracted is added to the concept index. By indexing study variables with ID pointers to concepts, Dug eliminates the need to calculate these groups dynamically, and eliminates redundant text stored across study variables mapping to the same ontological concept. Each indexed knowledge graph answer contains a JSON representation of the answer subgraph returned by TranQL and a pointer back to the ontological concept used in the original query.

### Search Engine

#### Search Functionality

Dug’s search API exposes three search endpoints for querying each underlying Elasticsearch index.

- **/search_var** - search for study variables matching a user’s query
- **/search_concepts** - search for ontological concepts matching a user’s query
- **/search_kg** - search for knowledge graph answers matching user’s query and an ontological concept id

Dug’s search API uses the default Elasticsearch algorithm (cosine similarity based on vector space model using TF-IDF weighting) (Elasticsearch, 2021) to rank and retrieve indexed metadata records. The search field weighting scheme prioritizes exact matches from the originally ingested text first, followed by synonyms added through annotation, and lastly related terms added through concept expansion.

#### Search User Interface

Dug’s user interface (UI) is a stand-alone React JS-based web application designed to provide an intuitive interface for navigating large collections of data. Dug’s minimalist UI design is intended to reduce the burden on users and empower them to discover search terms they are interested in exploring. Dug provides a search box that prioritizes exact phrase matching (AND logic) over partial matching (OR logic) (Fig. 5). Other features include auto-generated tabbing of search results by data type.

**Fig. 5:**
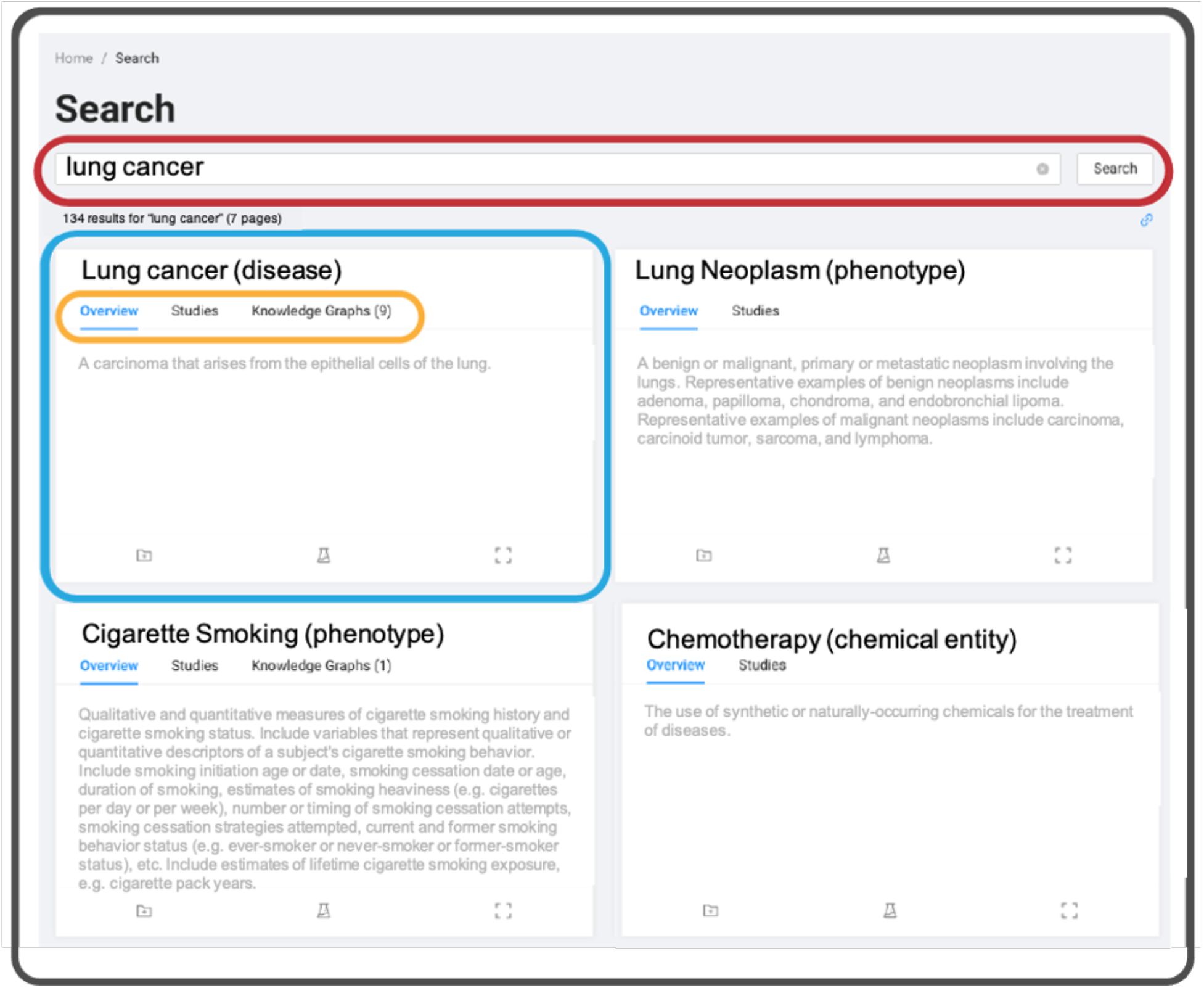
Dug’s UI aggregates search results for user queries (red) into higher-order ontological concepts (blue) based on NLP annotation. Links to knowledge graphs (orange) explain biological relationships between the query and each concept. Concepts may not contain knowledge graph links if synonymous with the search query (e.g., “lung cancer” vs. “adenocarcinoma of the lung”) or if TranQL did not return answers during concept expansion.

Dug’s UI organizes search results in two ways: by variable and by concept. When organized by variable, Dug returns study variables sorted by relevance, with each result containing information about the variable returned (e.g., parent study). When organized by concept (Fig. 5), Dug aggregates results into higher-level ontological concepts. Dug can then be used as a preliminary harmonization step to create *de novo* groups of similar variables based on NLP annotations (Fig. 5). By providing both approaches, Dug’s concept-based search gives users an exploratory look at the data landscape, while its variable-based search allows users to investigate specific variables of interest.

The defining feature of Dug’s UI is its ability to explain *why* it returns certain results (Fig. 6). When Dug returns a result based on text added during concept expansion, the UI fetches and renders the corresponding knowledge graph answer from the knowledge graph index to explain the connection (if available, answers include links to supporting peer-reviewed literature).

**Fig. 6:**
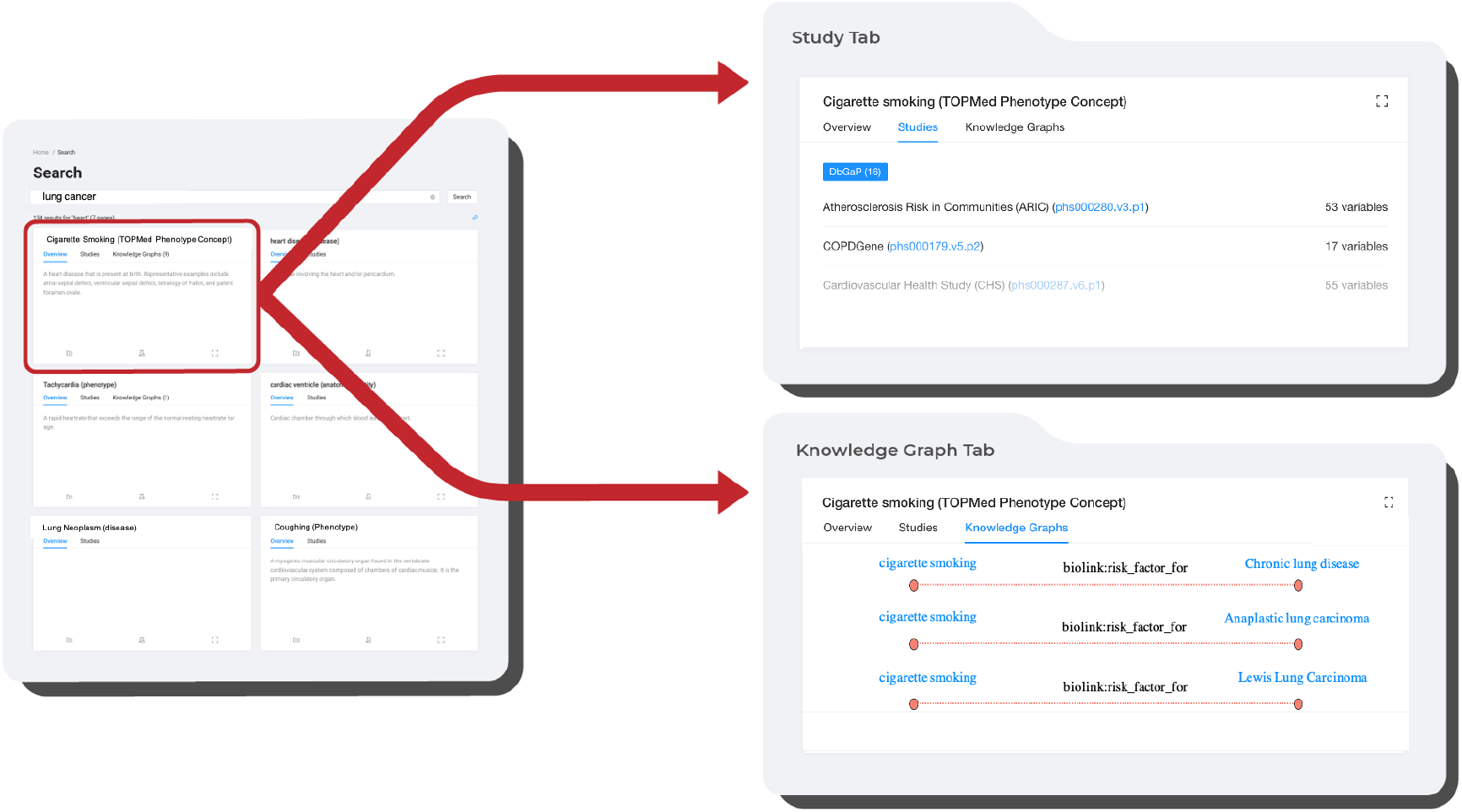
Dug results show datasets with variables relevant to a user’s query (e.g., “lung cancer” returns “cigarette smoking”), and knowledge graph disclosures for understanding why Dug considered those results relevant (e.g., cigarette smoking is a risk factor for lung cancer).

### Implementation

The Dug search app is implemented in Python 3.9 (https://python.org) and available via GitHub (https://github.com/helxplatform/dug). Dug packages core code and 3rd party services into a Dug container service deployable locally via docker-compose (https://docs.docker.com/compose/) or on Kubernetes (https://kubernetes.io/) clusters via a Dug helm chart. Along with Elasticsearch, Dug uses Redis (https://redis.io/) to cache API requests and minimize redundant calls to external services. Dug’s core services are externally configurable from a single file allowing users to specify annotation modules, external API endpoints, ontology normalization services, and query templates used during concept expansion (see Supplement). Dug’s ingest architecture leverages the Pluggy framework (https://pluggy.readthedocs.io/en/latest/) to define new metadata parser modules. Dug provides a makefile script to install the service locally. Lastly, all major commands can be invoked from the command line or via API calls to the service directly.

### Deployment on BioData Catalyst

We deployed Dug on the NHLBI’s BioData Catalyst ecosystem and indexed the TOPMed freeze 5b and 8 studies (excluding parent studies). Public data dictionaries downloaded directly from dbGaP were ingested and indexed for each of the 76 datasets included in these freezes.

15,911 of these variables were manually harmonized into 65 higher-order groups called “phenotype concepts” by data curation experts at the TOPMed Data Coordinating Center (DCC) (Stilp *et al.*, 2021). The TOPMed harmonized phenotypes were created to enable interoperability between TOPMed datasets by manually combining semantically similar dbGaP variables under a single term. To facilitate browsing, we annotated ingested variables with phenotype concepts as though they were external ontology identifiers so the underlying variables could be aggregated by these phenotype concepts in the web portal.

### Evaluation

We evaluated Dug’s performance against a purely lexical search strategy (default Elasticsearch; TF-IDF with “fuzzy matching”) to quantify the impact of Dug’s semantic awareness on recall and precision. We used the TOPMed phenotype concepts dataset as a framework for evaluation.

Variable descriptions were preprocessed prior to evaluation to characterize search performance more accurately. Steps included lowercasing descriptions, removing punctuation, removing the uninformative “Exam {integer}” pattern, normalizing whitespace, and de-duplicating to include only unique variable descriptions. We preprocessed the TOPMed Phenotype concept names by removing stop words (e.g., “the,” “in,” “of”), and expanding known abbreviations (e.g., “Resting arm systolic BP” becomes “Resting arm systolic BP blood pressure.”).

For each of the 65 TOPMed phenotype concepts, we queried concept titles against Dug’s indexed collection of dbGaP variables to see how well Dug could recapitulate the manually curated set of variables. Nearly 76% of the variables in each TOPMed phenotype concept contained a lexical match to the name of the concept we had used as a test query. To evaluate Dug’s performance on a less trivially simplistic dataset, we repeated the evaluation using synonyms of TOPMed phenotype concepts as test queries. We used synonyms from the Unified Medical Language System (UMLS) (Bodenreider, 2004) with no words in common with the original query; if no reasonable synonym could be chosen for a TOPMed phenotype, it was removed.

To quantify Dug’s performance, we used recall and precision. We report scores at the 10^th^ result (P@10, R@10), at the 50^th^ result (P@50, R@50) and at the n^th^ result (P, R) for every search. While (P, R)@10 is a standard metric, we chose (P, R)@50 as the maximum number of results a person would reasonably scroll through, assuming 25 results per page (Jansen and Spink, 2005).

Within this framework, each variable belonging to a given TOPMed phenotype concept represents a condition positive (P) we would expect Dug to return. True positive (TP) results were variables returned by the TOPMed phenotype concept to which they belonged. Conversely, false positives (FP) were variables returned by any TOPMed phenotype concept to which they did not belong. Recall and precision are calculated at results returned (10, 50, or n) for each TOPMed phenotype concept query.

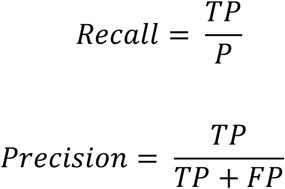

We evaluated Dug’s performance against a lexical search strategy as a baseline for how “findable” test datasets were without Dug’s semantic enrichment. We indexed the same set of study metadata into Elasticsearch using Dug’s ingestion pipeline but did *not* add any semantic annotations. Search results were retrieved using the same default Elasticsearch scoring algorithm used by Dug. This was done to eliminate variation between Dug and a baseline search due only to differences in the underlying lexical search strategies employed, allowing us to attribute differences in performance to Dug’s semantic annotation. There was no significance to using Elasticsearch beyond it providing a sophisticated lexical search strategy to make our evaluation results more directly interpretable.

A post-hoc analysis quantified the average semantic and lexical similarity between test queries and the top 50 results returned by Dug and Elasticsearch. Semantic similarity was measured using cosine similarity between test queries and search results after using the BioSentVec (Chen *et al.*, 2019) model to transform queries and phrases into sentence embeddings. The sent2vec (v0.2.2) Python package was used to load the pretrained BioSentVec model (Pagliardini *et al.*, 2018). Lexical similarity was measured using Levenshtein distance implemented in the python-Levenshtein Python package (v0.12.2).

To determine the statistical significance of differences in performance metrics between Dug and default Elasticsearch, we used nonparametric paired Wilcoxon signed rank tests to test the hypothesis that location shift for each performance metric was nonzero. We chose this nonparametric test as it allows violations of normality in the sample distribution of differences. Wilcoxon tests were performed using the Python SciPy (v1.7.3) package using two-sided alternative hypotheses and a critical value of 0.05 (Virtanen *et al.*, 2020).

## Results

The initial Dug deployment on the NHLBI Biodata Catalyst ecosystem successfully indexed 15,991 study variables from 76 genomics datasets. Dug augmented these study variables with 573 ontological concepts and 11,752 knowledge graph answers.

### Evaluation

The results of our initial evaluation (Supplemental Fig. 1) showed Dug outperformed default Elasticsearch in terms of mean recall (0.79 vs. 0.76, p < 0.001) but was not significantly different from Elasticsearch in terms of recall @ 10 and @ 50 (Supplemental Table 1). Additionally, Dug performed slightly but significantly worse than Elasticsearch in terms of precision @ 10 (p=0.022) and @ 50 (p=0.027). The biggest difference between Dug and a text-based method was Dug’s significantly lower total precision than Elasticsearch (0.07 vs. 0.28, p < 0.001), though this was expected as Dug is designed to return a larger set of more exploratory results.

After repeating the evaluation with synonyms of original queries (e.g., “lung adenocarcinoma” vs. “lung cancer”), Dug vastly outperformed lexical search. Shown in figure 7, Dug significantly outperformed default Elasticsearch in terms of mean recall (0.36 vs. 0.14, p < 0.001), recall @ 10 (0.08 vs. 0.04, p=0.033) and recall @ 50 (0.18 vs. 0.08, p=0.007). We found no significant difference between Dug and Elasticsearch in precision @ 10 (p=0.184) and precision @ 50 (p=0.223); Dug’s total precision was significantly worse (p=0.041), as to be expected with Dug’s exploratory results.

**Fig 7:**
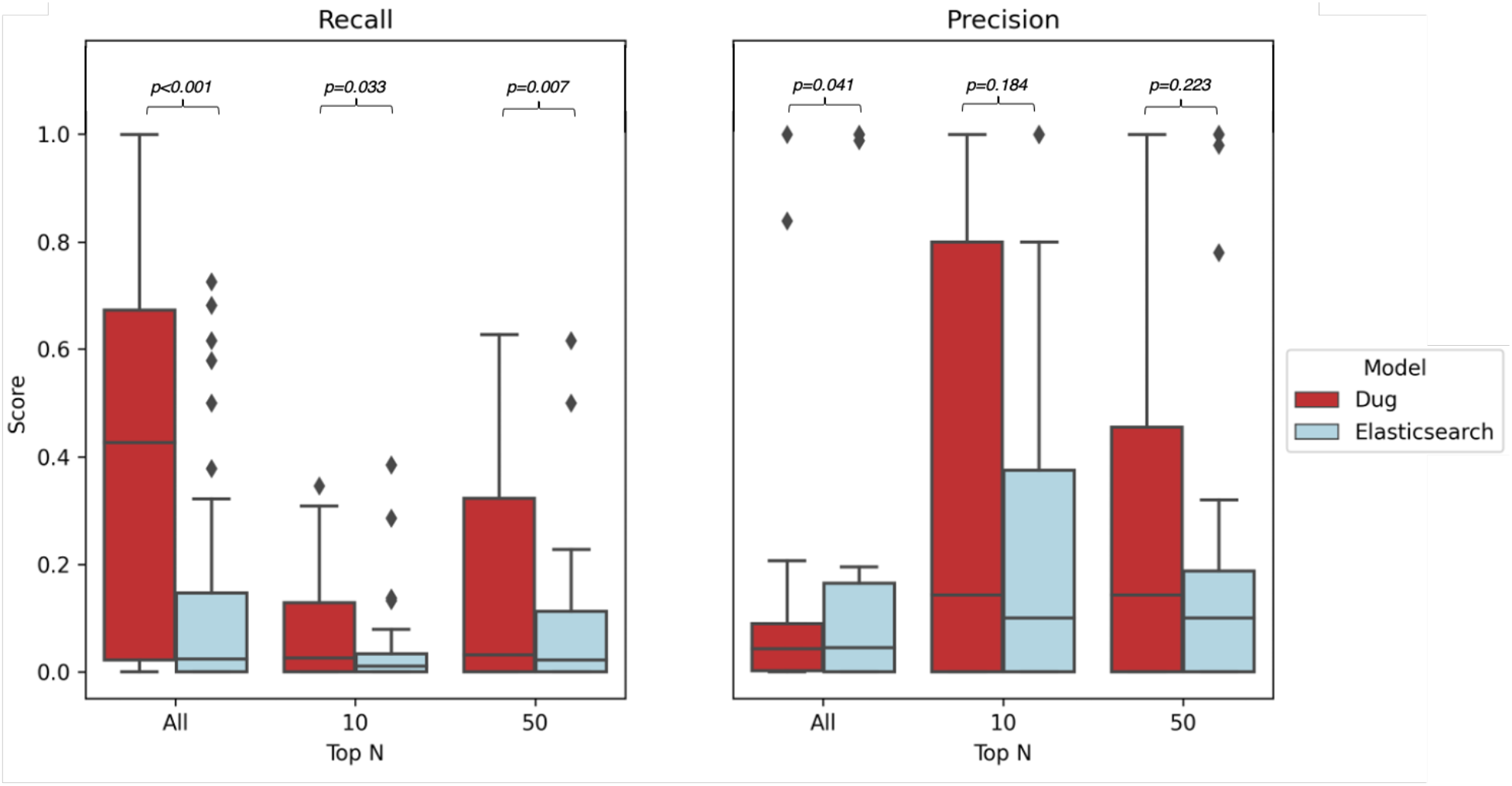
Dug vs lexical search performance using UMLS synonyms of TOPMed phenotype concepts to search for underlying variables. Left: Recall, R@10, R@50. Right: Precision, P@10, P@50. Dug significantly outperforms lexical search (implemented in Elasticsearch) in terms of total recall, R@10, and R@50. Precision @10 and P@50 were not significantly different between Dug and Elasticsearch. Though Dug’s total precision was significantly lower, this was expected given Dug’s objective to uncover exploratory connections.

Together, these results suggest Dug’s superior recall results from including semantically equivalent terms that don’t appear lexically similar to the query (e.g., “heart attack” vs. “myocardial infarction”). To test this hypothesis, we quantified the average semantic and lexical distance between queries and results returned by Dug and Elasticsearch. Dug’s results are on average significantly (p=0.021) more lexically divergent from the original query than Elasticsearch results, despite there being no significant difference (p=0.878) in the semantic distances of search results returned by each (Fig. 8). Together, these results suggest Dug outperforms the most sophisticated lexical search strategies when lexical diversity of the search space is high, as is typical for non-standardized biomedical datasets.

**Fig 8:**
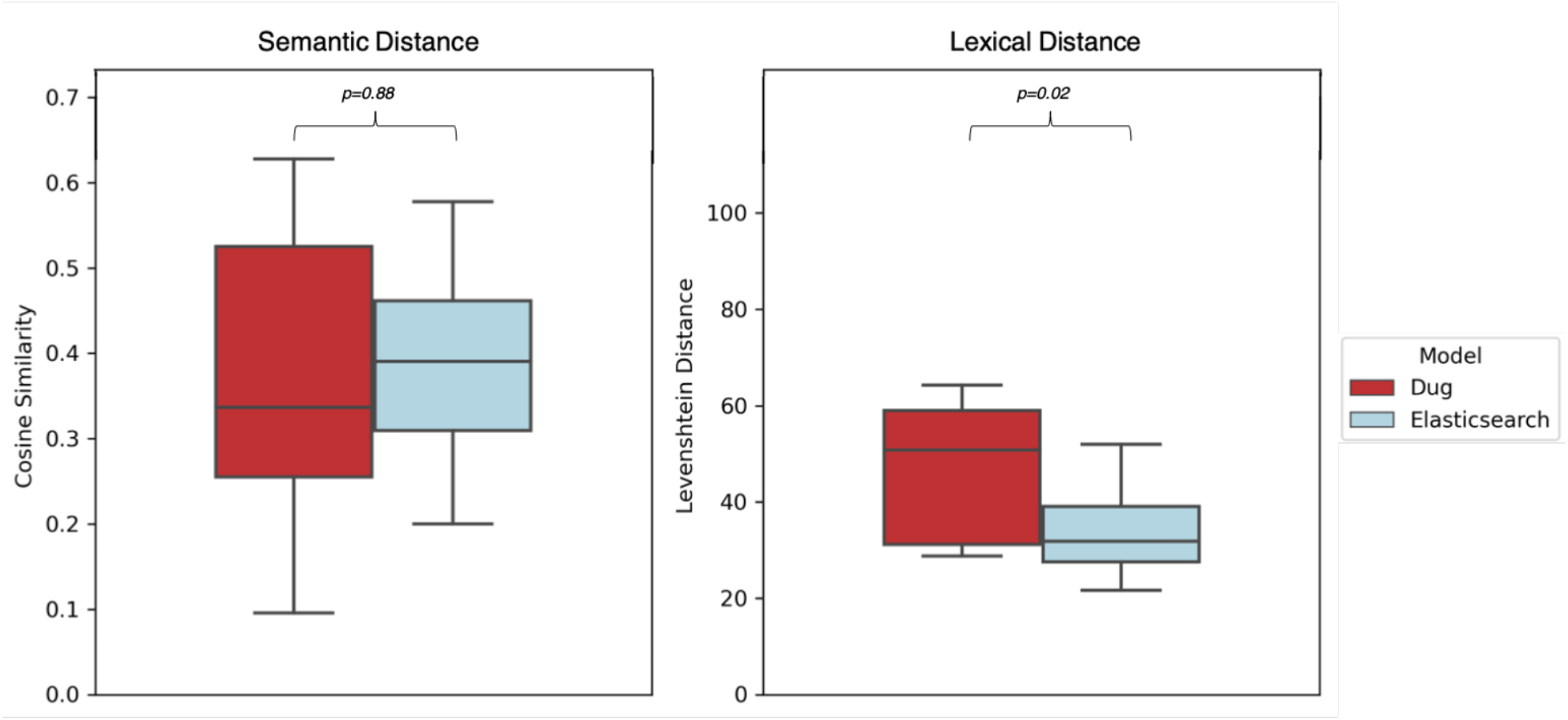
Semantic vs. Lexical distance of Dug and Elasticsearch search results. Left: Semantic distance between queries and top 50 results returned by each method. Semantic distance for each query is defined as the mean cosine similarity of the top 50 results in the pre-trained BioSentVec model vector space. Right: Lexical distance between each query and its top 50 results (mean Levenshtein distance). Dug and Elasticsearch return semantically similar results on average (p=0.88), yet Dug’s results are significantly more lexically divergent from the query (p=0.02) with no loss in precision.

## Discussion

The exponential increase in publicly available datasets has created a need for comprehensive search tools that identify datasets relevant to a researcher’s particular scientific question. To help researchers better navigate this new data landscape, we created Dug: a semantic search tool for biomedical datasets leveraging ontological knowledge graphs to intelligently suggest relevant connections derived from peer-reviewed research.

Our results demonstrate Dug’s ability to find relevant datasets regardless of lexical expression of a query. When compared to a sophisticated lexical search strategy like Elasticsearch, Dug returns more relevant results with no significant difference in the precision of the top 10 or top 50 results. Dug also outperforms lexical search particularly when the search space is characterized by the type of high lexical diversity typical of biomedical data repositories.

Dug’s potential for data exploration and discovery is perhaps its greatest and least quantifiable strength. Dug’s ability to surface relevant datasets may be most useful when the user is not sure what they are looking for. For example, the relevance of “asbestos exposure” variables to a search for “lung cancer” datasets is contextual and subjective, making it difficult to quantify when and how second and third order connections might be useful. A key innovation of Dug is its ability to provide exploratory connections without sacrificing the precision of top results.

Dug’s modular design and stand-alone companion web portal can flexibly fit myriad use cases. For example, a centrally hosted version of Dug could index multiple data repositories to service a much larger user base. Additionally, smaller data coordinating centers like NIDDK central repository (Rasooly *et al.*, 2015; Cuticchia *et al.*, 2006) or large data ecosystem initiatives like NIH’s HEAL Data Ecosystem (U.S. Department of Health and Human Services) could use Dug to search across non-standardized data from diverse consortium members via a single portal without the need for significant manual curation.

Future work focuses on known limitations and responding to user feedback from the BioData Catalyst consortium. A principal concern is parallelizing the indexing process to index multiple datasets simultaneously and increase the throughput for larger datasets. Additionally, we are evaluating various strategies for improving ranking search results by relevance based on input from current users. We believe Dug provides a powerful, flexible tool for searching intuitively across complex data resources that are increasingly common in the biomedical data landscape.

## Acknowledgements

The authors would like to thank Ben Heavner, Adrienne Stilp, and Ingrid Borecki for their invaluable feedback during development. We also want to thank members of the BioData Catalyst Fellows program for feedback and testing.

