## Supplemental Material for "Dug: A Semantic Search Engine Leveraging Peer-Reviewed Knowledge to Span Biomedical Data Repositories"

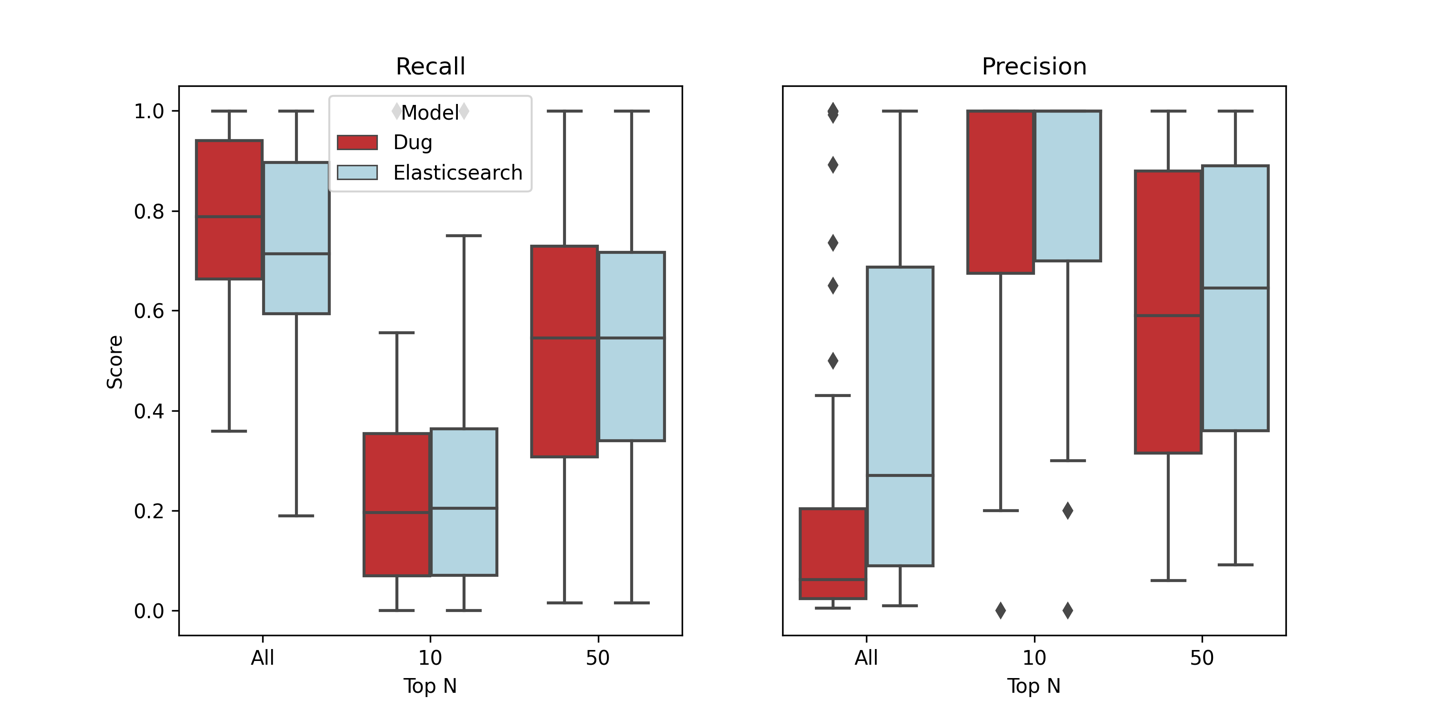
*Supplemental Fig. 1. Dug vs. Elasticsearch recall and precision when using TOPMed phenotype concept names as search queries. These metrics capture Dug and Elasticsearch performance on a simplistic test dataset where 76% of search results contain a lexical match to the search query. Under these conditions, Dug’s total recall is slightly but statistically significantly higher than Elasticsearch. Though Dug performs significantly worse in terms of total precision, this is to be expected as Dug is designed specifically to return exploratory connections. Dug performs slightly worse in terms of precision @ 10 and precision @ 50. These results demonstrate that even under more ideal circumstances for a lexical search strategy, Dug imposes minimal tradeoffs.*

Supplemental Table 1. Significance testing results for comparisons shown in figures. Here we show the results of paired Wilcoxon rank tests for significant differences between Dug and Elasticsearch across various performance metrics. The Query Group column represents whether Phenotype concept names or Synonyms were used as queries for computing recall and precision.

| ***Query Group*** | ***Metric*** | ***Top N*** | ***p-value*** |
| --- | --- | --- | --- |
| *Phenotype Concept Names* | *Recall* | *All* | ***<0.001*** |
| *Phenotype Concept Names* | *Recall* | *10* | *0.070* |
| *Phenotype Concept Names* | *Recall* | *50* | *0.664* |
| *Phenotype Concept Names* | *Precision* | *All* | ***<0.001*** |
| *Phenotype Concept Names* | *Precision* | *10* | ***0.022*** |
| *Phenotype Concept Names* | *Precision* | *50* | ***0.027*** |
| *Synonyms* | *Recall* | *All* | ***<0.001*** |
| *Synonyms* | *Recall* | *10* | ***0.033*** |
| *Synonyms* | *Recall* | *50* | ***0.007*** |
| *Synonyms* | *Precision* | *All* | ***0.040*** |
| *Synonyms* | *Precision* | *10* | *0.184* |
| *Synonyms* | *Precision* | *50* | *0.223* |

### Dug Common Metadata Format

The first step of Dug ingestion and indexing is to parse study variables into a list of *DugElement* objects. The required fields for a *DugElement* are shown in Supplemental Table 2.

*Supplemental Table 2. Names and descriptions of fields in DugElement metadata model.*

| **Field** | **Description** | **Example** |
| --- | --- | --- |
| element_id | Unique id to identify a study element (e.g., variable id) within a collection (e.g., study, image series) | phv00082951.v1.p3 |
| element_name | Element name that will be displayed on UI (e.g., variable name) | LDL in blood |
| element_description | Text description of element used as input for variable annotation. | “Cholesterol (LDL) measure at first visit” |
| element_action | External link or action to more info about element | www.ncbi.nlm.nih.gov/projects/gap/cgi-bin/variable.cgi?study_id=phs000209.v13.p3&phv=00082951 |
| data_type | Element data type or source repository | DbGaP |
| collection_id | Unique id of containing study or collection (e.g., study id) | phs000209.v13.p3 |
| collection_name | Name of containing study or collection | Multi-Ethnic Study of Atherosclerosis (MESA) |
| collection_action | External link or action to more info about the containing collection | www.ncbi.nlm.nih.gov/projects/gap/cgi-bin/study.cgi?study_id=phs000209.v13.p3 |

In short, a *DugElement* is designed to make artifacts associated with a particular scientific study searchable through the Dug interface. Unlike more sophisticated metadata models like DATS, a *DugElement* makes few assumptions about the structure of the data being indexed; this design decision was explicitly chosen to flexibly accommodate a broad range of scientific data types without the need to require additional metadata fields: study variables, images, or even applications or software tools can all be represented as *DugElements*. Also, unlike many dataset search engines that collect *only* study level metadata, Dug actually does not index *any* study level metadata. As such, a study/collection is made findable because its underlying artifacts are findable and can link back to a parent/containing study. The Dug UI utilizes the *collection_id* field to aggregate search results by study so that it appears Dug is searching at the study level.

### Extending Dug to index data from new sources

When extending Dug to support indexing a new input data type (e.g., DbGaP dictionaries), one needs to define how elements from that input type can be parsed into the *DugElement* structure. Typically, this means being able to parse study data dictionaries from various sources and map individual columns to a corresponding *DugElement* field. In Supplemental Table 3, we show a simple example of a csv data dictionary that could be converted to a list of *DugElements*.

*Supplemental Table 3. Example data dictionary describing experimental variables measured by a study.*

| **Study** | **Study_id** | **Study_link** | **Var_ID** | **Var_Name** | **Var_Desc** | **Var_Link** |
| --- | --- | --- | --- | --- | --- | --- |
| My Heart Study | myrepo.myheartstudy | myrepo.myheartstudy.com | myvar_1 | HRATK | Status of whether patient has had a myocardial infarction in last 6 months | myrepo.com/mydataset/myvar_1 |
| My Heart Study | myrepo.myheartstudy | myrepo.myheartstudy.com | myvar_2 | STAT | Was patient taking a statin drug at any point in the past 6 months | myrepo.com/mydataset/myvar_2 |
| My Heart Study | myrepo.myheartstudy | myrepo.myheartstudy.com | myvar_3 | LDL | LDL cholesterol measure at most recent clinic visist | myrepo.com/mydataset/myvar_3 |

Assuming all data dictionaries from the fictional *myrepo* repository have this structure, we could index all *myrepo* studies by creating a parser class in Dug that defines a method to map these variables to *DugElements*. A pseudocode example is shown in Supplemental Figure 2.

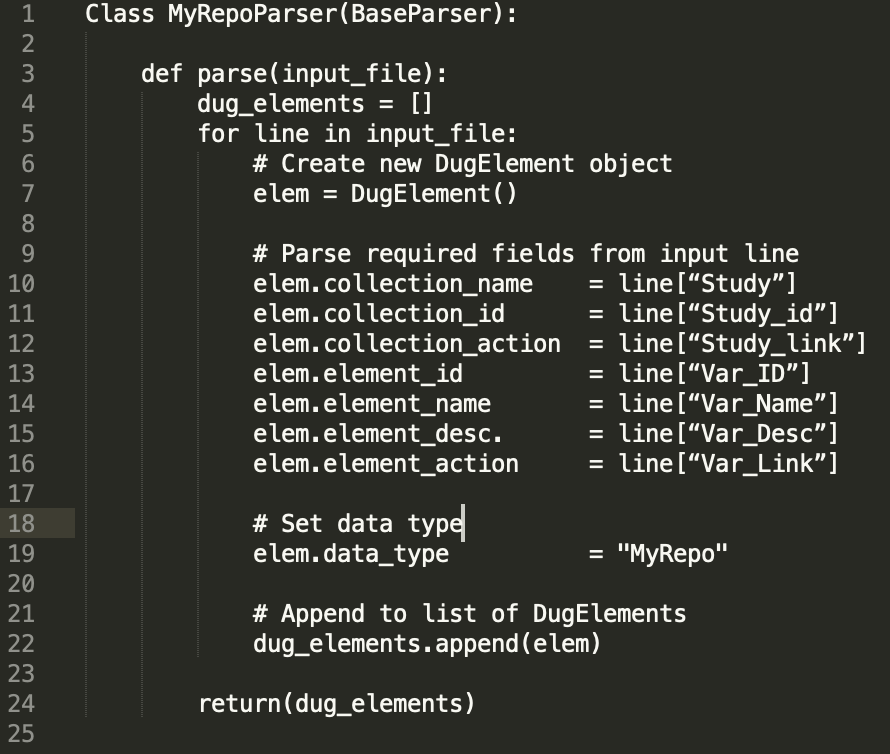

*Supplemental Figure 2. Pseudocode defining a parser class to map elements from the csv in Supplemental Table 3 to a list of DugElement objects that can be indexed by Dug.*

Because Dug allows the user to specify the parser on the command line, Supplemental Fig. 3 shows how the Dug crawl command could then be invoked to index *any* study from the *myrepo* repository.

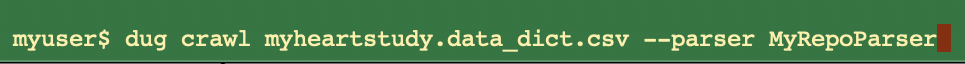

*Supplemental Fig. 3. Command line invocation to index a myrepo study using the parser defined above.*

### Configuring Additional Dug Options

Many aspects of the Dug indexing process can also be configured via an external configuration file. Here, we discuss the impact of these configuration options on Dug’s behavior.

Most consequentially, Dug allows the user to configure *all* of the external API services it uses to swap out components as users see fit. Shown in supplemental Figure 3, the user can configure which API services are used for named-entity recognition, normalization, synonym finding, and fetching additional ontology information. These have the potential to drastically alter Dug’s runtime behavior based on the sensitivity and accuracy of the underlying tools. Most importantly, this means Dug can easily be updated to utilize emerging or state of the art ML annotation tools without ever touching any code.

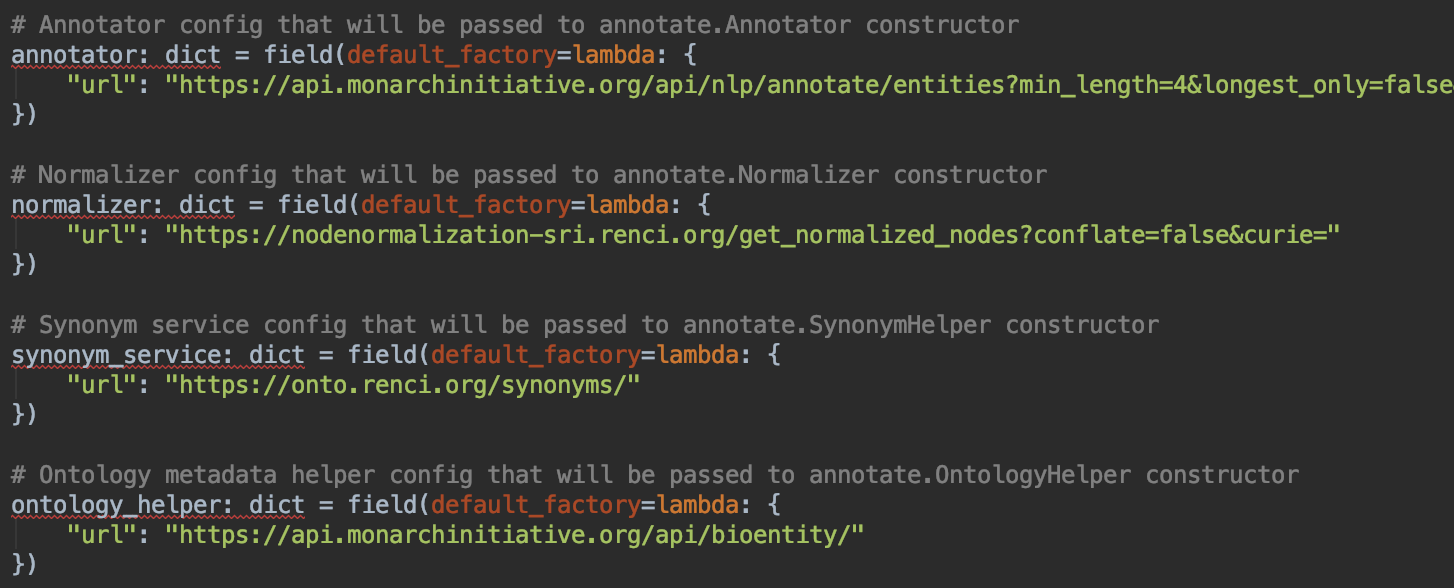

*Supplemental Figure 3. Dug’s configuration module allows users to easily change which API services are used for named entity recognition and annotation.*

Additionally, Dug allows users to manually specify a list of stop words, as well as mappings used to expand common abbreviations (Supplemental Fig. 4). During indexing, Dug’s preprocessing module removes stop words and expands any detected abbreviations into their long forms prior to submitting a *DugElement* for annotation. As shown in Supplemental Fig. 4, this behavior is easily externally configurable.

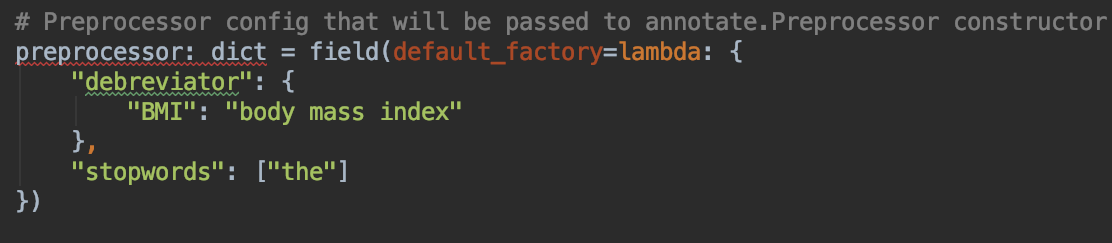

*Supplemental Figure 4. Configuring Dug’s debreviation and stopword behavior can aid the success of annotation steps.*

Finally, Dug also allows the user to define the TranQL queries that will be used during concept expansion.

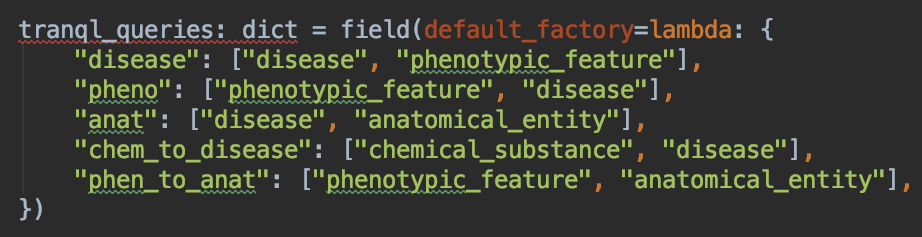

*Supplemental Figure 5. Dug’s configuration module allows users to define the TranQL queries that will be used during concept expansion to gather related biological concepts.*

In the example shown in Supplemental Figure 5, each entry in the map defines a one-hop query. For example, the statement [“disease”, “phenotypic_feature”] is interpreted by Dug as the following TranQL query template:

FIND disease -> phenotypic_feature WHERE phenotypic_feature == {query_ontology_id}

In English, when a *DugElement* is annotated as an ontology identifier recognized by TranQL as a phenotypic feature (e.g., dry cough, nearsightedness), TranQL will fetch ontology identifiers of diseases characterized by those phenotypic features (e.g., emphysema, albinism).
